# Environment shapes the association between the skin microbiome and infection dynamics of the Dermocystid pathogen *Amphibiothecum* in Palmate newts (*Lissotriton helveticus*)

**DOI:** 10.1101/2025.11.26.690651

**Authors:** Xavier A Harrison, Emma Remotti, Frances C Clare, Kevin Hopkins, Thalassa McMurdo Hamilton, Kirsten McMillan, Charlotte L Clarke, Tiffany Tsui, Anna Meredith, Trenton WJ Garner

## Abstract

Infectious diseases are a leading cause of biodiversity declines, but the factors shaping the prevalence and intensity of wildlife disease outbreaks remain unresolved. Host-associated microbes are likely a key component of host resistance to infection, but a host’s ability to recruit such protective microbes may be restricted by the availability of microbes present in their immediate environment. Here we investigate the associations between the abiotic environment, skin microbiome, and infection with the Dermocystid pathogen *Amphibiothecum meredithae* in Palmate newts (*Lissotriton helveticus*) on the Isle of Rum, Scotland. We found more acidic ponds to be associated with higher skin microbiome diversity and lower pathogen prevalence. Predicted functional analysis of microbiota composition identified bacterially-mediated pathways linked to host immunity upregulated in skin microbiotas associated with more acidic ponds and lower infection prevalence. This study highlights the potential importance of the environment in modulating host- microbiome-pathogen relationships and patterns of infection in the wild.

## INTRODUCTION

Microbial communities that live on and in wildlife are increasingly recognized as key determinants of wildlife health (Worsley et al., 2024). Microbes can be recruited by hosts through both environmental (indirect) and social (direct) transmission pathways (e.g. Raulo et al., 2024), but variation in the environment also shapes what microbes are available for both forms of colonisation. Abiotic variables can alter microbial persistence in the environmental microbiota, and therefore their availability (Dupont et al., 2016; Glaser et al., 2015) and stability (Buttimer et al., 2024), as well as alter pathogen persistence in the environment and therefore infection risk (e.g. Turner et al., 2021). Thus, spatial and temporal variation in patterns of infectious disease may be driven by factors other than host immunity, and disease dynamics may result from ‘environmental mismatches’ that are suboptimal for both maximising recruitment of protective microbes and minimising pathogen prevalence. Much of this has been determined through controlled laboratory studies that have revealed evidence of environmental effects on host-pathogen (Ellison et al., 2020; Garner et al., 2011; Padfield et al., 2020) and pathogen-microbiome (Muletz-Wolz et al., 2017) interactions. Comparable studies from complex natural systems are rare (but see Buttimer et al., 2024). Integrating data on hosts, pathogens, microbiotas and the environment in which these three agents interact is crucial for understanding contemporary patterns of disease prevalence and severity (Bernardo-Cravo et al., 2020), as well as predicting future risks of disease outbreaks due to climatic extremes (e.g. Hector et al., 2022).

Amphibian parasites from the order Dermocystida (class Mesomycetozoa), a clade at the intersection between fungi and animals (Mendoza et al., 2002; Rowley et al., 2013), are pathogens of amphibians that can cause host population demographic declines (Feldman et al., 2005; Raffel et al., 2008). Infected individuals display subcutaneous cysts, ulcerative lesions, and oedema, and may die as a result (González-Hernández et al., 2010; Raffel et al., 2008). Dermocystida have an environmental infectious stage (endospores) through which they infect hosts in freshwater habitats (Fiegna et al., 2017; Gleason et al., 2014; Woodhams et al., 2023). One of these hosts is the palmate newt (*Lissotriton helveticus*), recently described suffering from visible infections with the dermocystid *A. meredithae* in France and Scotland (Fiegna et al., 2017; González-Hernández et al., 2010). On the Isle of Rum in Scotland, where *L. helveticus* is the only amphibian species, newts at a subset of breeding sites presented infections that appeared far more severe than those affecting palmate newts in France; these cases involved ulcerative lesions, subcutaneous haemorrhages and extensive skin oedema that impaired the development of granular glands (Fiegna et al., 2017). This led the authors to speculate that *A. meredithae* might disrupt gland function and antimicrobial peptide production, which in turn might impact maintenance of a healthy skin microbiota. In support of this, a study of *Amphibiocystidium* infection and microbiome composition in Italian stream frogs (*Rana italica*) found that infection was associated with a less variable but distinct bacterial microbiota composition (Federici et al., 2015).

Patterns of decreased microbial diversity and both increased prevalence and severity of infections have been reported for amphibian hosts and pathogens, but the mechanism behind these patterns is often uncertain. What is clear, though, is that more severe infections and/or increased prevalence are often associated with environmental conditions that differ from locations where amphibian hosts appear healthier (Bosch et al., 2018; Clare et al., 2016; Garner et al., 2011; McMillan et al., 2020; Price et al., 2019; Spitzen-Van Der Sluijs et al., 2014). Palmate newts are known for their ability to occupy a diversity of environments, as their distribution ranges from Scotland to the Mediterranean. Temperature varies significantly across their distribution, but Palmate newts are one of a variety of amphibian species noted for their ability to tolerate relatively low pH (Brady & Griffiths, 1995; Griffiths et al., 1993; Pierce, 1985; Pottier et al., 2022). However, low pH ponds are known not to provide the optimum environment for growth and development (Brady & Griffiths, 1995; Griffiths et al., 1993). Rum is the remnant of a volcanic crater dominated by heath, grassland and blanket bog. While more permanent water bodies can trend towards neutral pH, run-off from peat soils generate less persistent water bodies that predominate the landscape (Lowe, 1998). pH values as low as 4.38 have been reported in these impermanent water bodies (Fryer & Forshaw, 1979). The landscape of Rum, therefore, proffers a range of environmental conditions across which the interaction between newts, *A. meredithae* and patterns of recruitment of environmental microbes can be investigated.

We tested the hypotheses that abiotic traits such as water pH and temperature are associated with both differential skin bacterial microbiota composition and infection prevalence of the dermocystid parasite *Amphibiothecum meredithae* in Palmate newts on Rum. First, we used repeat surveys of 21 ponds on the Isle of Rum to determine spatial variation in *Amphibiothecum* infection prevalence and mortality, as well as associations with abiotic traits like pH and water temperature (Figure 1). In a subsequent visit, we swabbed newts from 4 of these sites known to differ in infection prevalence and abiotic environment to examine the links between environment, bacterial skin microbiota, and disease. We tested the prediction that within sites, infection with *Amphibiothecum* will result in decreased alpha diversity and differential community composition (beta diversity). To disentangle direct effects of the abiotic environment on the pathogen from indirect effects via the microbiome, we use predicted functional analysis of the bacterial microbiota from infected and uninfected newts to identify putative functional mechanisms through which differences in microbiome may be linked to host immunity.

**Figure 1:**
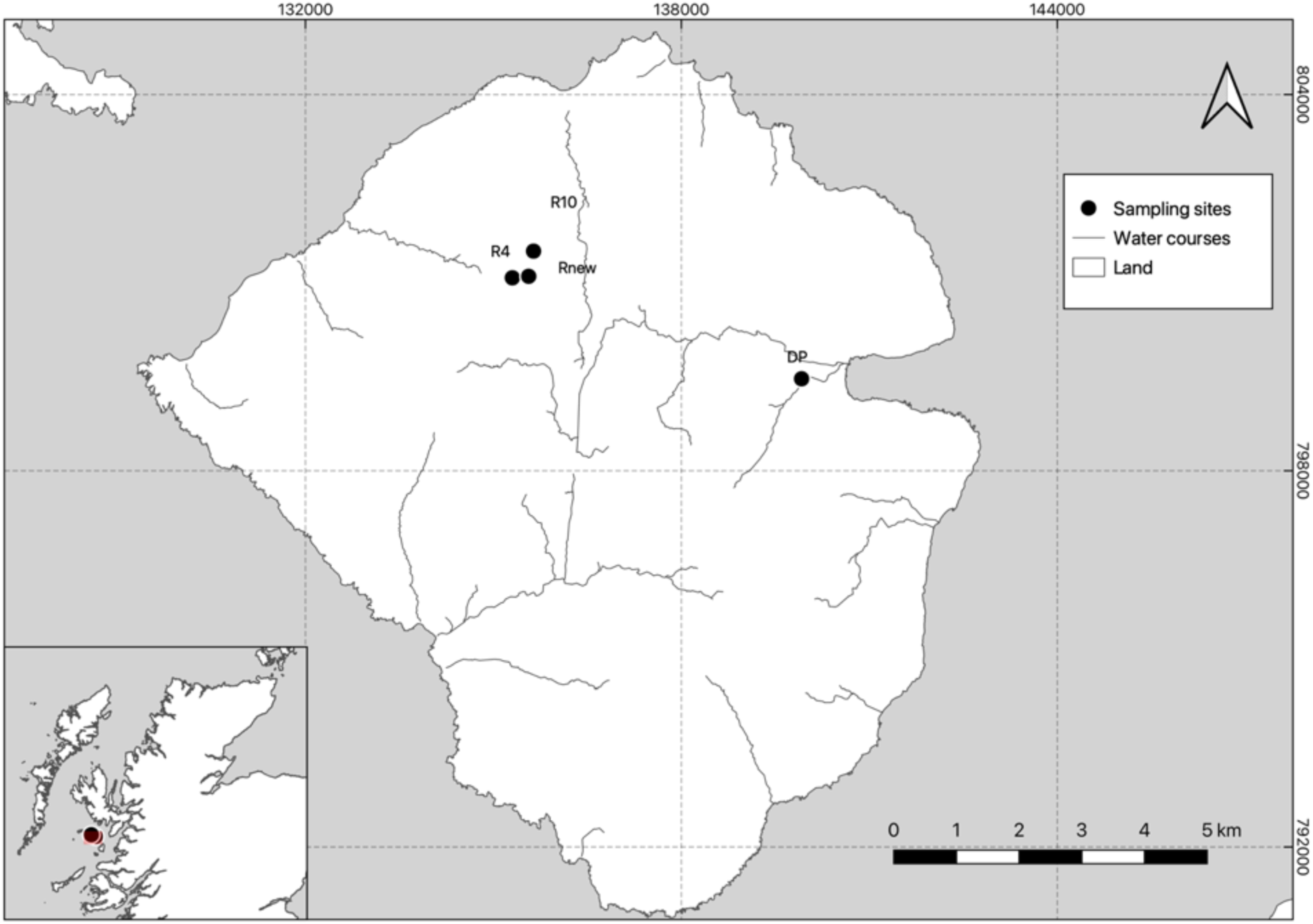
Map of the Isle of Rum, Scotland (N 57°00ʹ55.1ʹʹ, W 006°16ʹ53.3ʹʹ), showing locations where Palmate Newt (*Lissotriton helveticus*) skin microbiome samples were collected. Signs of *Amphibiothecum meredithae* infection were detected at 2 out of 4 sites (R4 and Rnew).

## METHODS

### Ethics Statement

All sampling and capture protocols were reviewed and approved by the ZSL Animal Ethics committee before work commenced. Of particular focus was the effects of handling and swabbing on the skin of visibly diseased animals. Our previous work (Fiegna et al., 2017) served as evidence that the work could be done without further compromising health of diseased animals.

### Amphibiothecum prevalence and skin microbiome Surveys

The Palmate newt (*Lissotriton helveticus*) is widely distributed across the Isle of Rum, a small island in the Inner Hebrides, Scotland. Most of the island is protected by Scottish Natural Heritage and has been classified as a National Nature Reserve since 1947. Vegetation types in the study areas were predominantly Blanket bog, Wet heath and *Calluna* heath. Palmate newts are the only amphibian species present on the island, inhabiting the numerous dystrophic lakes disseminated across the landscape.

Surveys to establish baseline prevalence of infection and correlates of infection took place during May 2012 to coincide with the breeding season, which is from approximately February until July (McMurdo Hamilton, 2012). We selected 20 ponds where high infection prevalence had previously been recorded. We recorded water temperature, altitude and pH of each pond three times per visit and at different parts of the pond using a HANNA meter.

Palmate newt microbiome samples were collected in May 2015 from 4 different pond sites (Figure 1). Two ponds (R4 and Rnew) were selected as locations where infections had been previously characterized, while the remaining two sites were selected as control, as no visible signs of infection had been detected over the course of the sampling season (Clarke, 2017). To collect skin microbiome samples, animals were dip netted and placed into large tubs of pond water. Wearing disposable gloves, each newt was then briefly rinsed with sterile water and sampled by gently rubbing the dorsal and ventral surface of the body with cotton swabs. Swabs were immediately put on ice and kept at -20°C until the end of the sampling season, then stored at -70°C until sequencing. Before release, all newts were sexed, measured for snout to vent length (SVL) and inspected for any visible signs of infection, recording severity of visible *Amphibiothecum* symptoms according to the criteria illustrated by Fiegna et al., (2017). Briefly, healthy newts were assigned a code of ‘0’, newts with small cysts were classified as category ‘A’, simple cysts with nodular swellings were type ‘B’, clustered swellings and cysts sometimes associated with discolouration and skin lesions were classed as type ‘C’. Lastly, individuals with widespread subcutaneous oedema were classed in category ‘D’. As only 2 individuals showed lesions of type ‘D’, category ‘C’ and ‘D’ were merged (category ‘C/D’) for analysis purposes. Environmental variables including water pH, air and water temperature were also recorded at each site.

### 16S Sequencing, Bioinformatics and Statistical Analyses

NGS libraries were prepared by targeting the v4 region of the 16S rRNA gene using an adapted protocol from Kozich et al. (2013), as described by Harrison et al. (2019). Negative controls were included during library preparation to help determine levels of background sample contamination. We also sequenced mock communities (positive controls) to allow us to benchmark diversity estimates. Samples were loaded on v2 chemistry 500 cycle cartridges for sequencing using 250 bp reads on an Illumina Miseq. All bioinformatics steps and statistical analyses were carried out in the R software, version 4.2.0 (R Core Team, 2024). The *dada2* pipeline (Callahan et al., 2016) was implemented using standard parameters as illustrated in the online tutorial to filter and trim 16S raw reads, remove chimeras, assign taxonomy and estimate error rates. Subsequent read processing and removal of mock and negative sequences was carried out using *phyloseq* (McMurdie & Holmes, 2013). Potential contaminants were then excluded from the dataset using the *decontam* package (v 1.16.0; (Davis et al., 2018), using negative controls as a reference and specifying a probability threshold of 0.5. Using these criteria, 33 out of 13903 total bacterial amplicon sequence variants (ASVs) that matched negative controls were removed from the dataset. To remove bias caused by low-abundance sequences, taxa with fewer than 100 reads were removed from the dataset, leaving a total of 4420963 reads from 1217 ASVs. Read depth ranged from 10542 to 83568 reads per sample, with a mean read depth of 36146.27.

The full reproducible workflow for analysis is included as an RMarkdown document on GitHub (see Data Accessibility), and so we provide a summary of statistical approaches here. Alpha Diversity: We rarefied sequencing data to the lowest sequencing depth of 10542 using a randomly generated seed and calculate d alpha diversity as observed richness for all samples. We modelled the relationship between alpha diversity, infection and sampling site using a generalised linear model with a Negative Binomial errors, with observed richness as a response variable and site, infection code (0, A, B or C/D), sex and mean-centred SVL as predictors. We did not include site as a random effect, as fitting a random effect with less than five levels can render inaccurate variance estimations (Harrison *et al.,* 2018). We used the R package *brms* (Bürkner, 2018, 2021) to fit models and assessed adequacy of fit using posterior predictive checks. We used full model tests (Forstmeier & Schielzeth, 2011) to control Type I error rates, and LOO-IC (Vehtari et al., 2017) to select among models, where a Δ6 LOO-IC score indicates strong support for the full model.

Beta Diversity: To quantify differences in community structure between sites and infection codes, we ran a principal component analysis (PCA) on the distance of the centred log-ratio transformed (CLR) data (also Aitchinson distances). Given the compositional nature of microbiome datasets, using a clr transformation rather than rarefaction prevents loss of data which can affect the relative abundances of individual taxa and consequently their position in the ordination space (Gloor *et al.,* 2016). We used PERMANOVA and CCA models fitted in *vegan* (v2.6-2; Oksanen et al., 2025) to test the effects of site, pH and infection code as predictors.

Differential Abundance and Predicted Functional Analyses: Differences in abundance of bacterial phyla across sites and infection statuses were visually inspected through stacked barplots of the top 5 most abundant phyla across all samples. We used General Linear Latent Variable Models (GLLVMs) to quantify differential abundance of bacterial genera using the R package *gllvm (*Niku et al., 2019, 2024). We used the software MicFunPred (Mongad et al., 2021) to predict functional profiles from our 16S rRNA bacterial community sequence data. We used the R package *mixOmics* (Rohart et al., 2017) to fit sPLS-DA models to identify variation among sites and infection classes in predicted microbiome function. We also used *mixOmics* to generate ordination and cluster image maps.

## RESULTS

### More Acidic Ponds Are Associated with Lower Amphibiothecum Prevalence

In the 2012 field surveys, we sampled 579 newts at 20 sites, spread over 59 visits. Average prevalence of infection across sites ranged from 5.3 – 80.6% (range 0-100% for individual site visits; Fig. 2A). Using Bayesian Mixed Effects Modelling, we detected a positive relationship between pH and infection prevalence (Fig 2A; Negative Binomial GLMM; effect of pH 0.68, 95% credible intervals [0.18 – 1.18]). Ponds with higher disease prevalence (⩾50%) seemed to be around physiological pH, while newts sampled at low pH ponds (<5.5) exhibited an average prevalence of infection of approximately 20%. Sites from 2012 with higher prevalence of infection also exhibited higher average mortality when controlling for sampling effort (Fig. 2B). pH and temperature exhibited a positive association when controlling for repeated measures at sites and temporal effects (Fig. S1; Table S1A). We found no association between altitude and pH (Figure S2; Table S1B).

**Figure 2.**
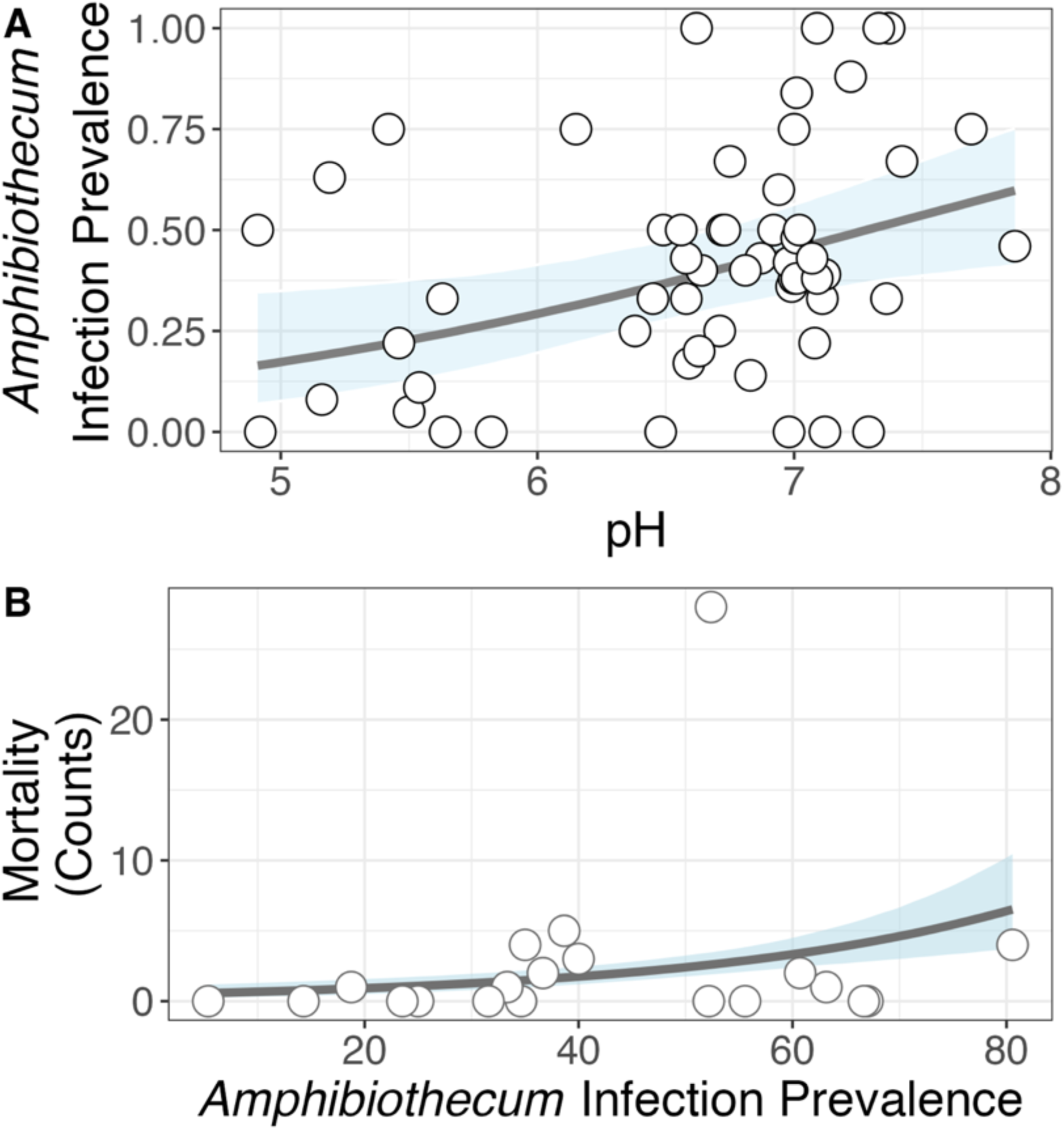
Infection Prevalence and Mortality Data from the 2012 Surveys. **A**) More acidic ponds (lower pH) are associated with lower infection prevalence of *Amphibiothecum.* Points are raw data from 20 sites surveyed multiple times (n=56 visits where 1 or more newts were sampled). Line and shaded area are mean and 95% credible intervals of a Bayesian general linear mixed model controlling for site effects. Visits where no newts were caught were excluded from this analysis. **B)** Site with higher average prevalence of infection also exhibit higher frequencies of dead (infected) newts. Points are average data per site over multiple surveys (prevalence) by total per-site mortality. Line and shaded area are mean and 95% credible intervals from a Bayesian general linear model with Poisson errors, controlling for survey effort.

### Skin Bacterial Community Composition Varies by pH and Infection

Infection prevalence for the 4 sites sampled in 2014 was similar to values from 2012, except for one site (DP) from which we recovered no infected newts (Table 1). We detected marked variation among these 4 sites in mean bacterial richness, measured as number of observed ASVs (Figure 3A; Table S2). There was no evidence that bacterial richness varied by sex or size (snout-vent length; Table S2). This model explained 74% [95% credible interval 65-79%] of variation in bacterial richness. Further modelling revealed a non-linear relationship between pH and bacterial richness, where more acidic ponds supported more diverse skin microbiota (Fig. 3B; Table S3). We also detected a negative influence of *Amphibiothecum* infection on richness (Table S3). Exploring this pattern in the two sites where prevalence was high (R4 & Rnew) revealed a significant infection:site interaction (Table S4) where only in R4 was alpha diversity reduced in infected newts (Fig. 3C).

**Figure 3.**
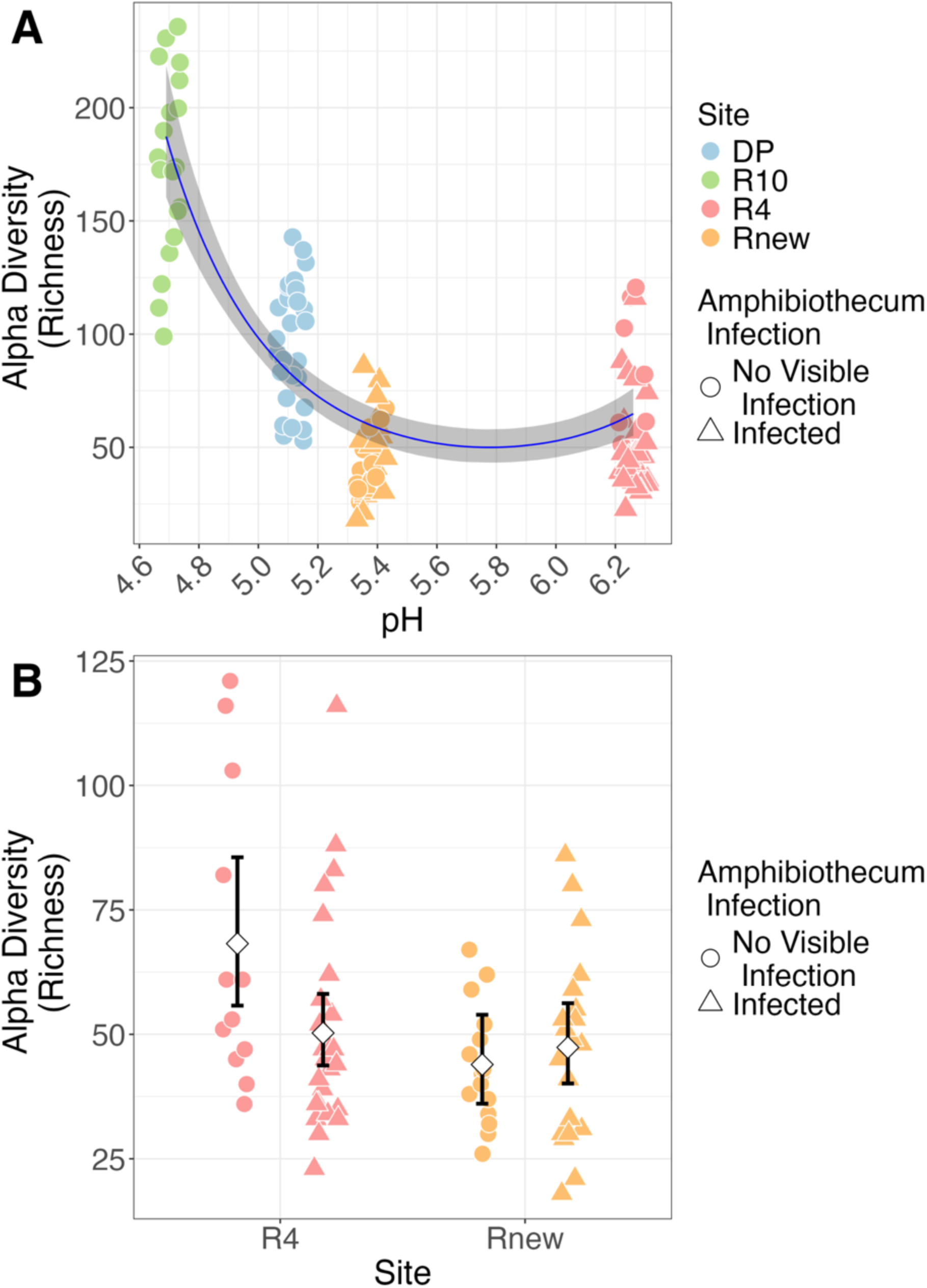
Palmate newt bacterial richness varies by site, pH and infection (A) Lower pH ponds are associated with greater skin bacterial richness. (B) Differences in bacterial richness by infection category within sites, where only R4 demonstrated a significant reduction in richness for infected newts.

**Table 1.** Prevalence of visible signs of infection with *Amphibiothecum meredithae* in Palmate newts across four sites on the Isle of Rum, Scotland used for microbiome profiling. Mean prevalence per site is calculated over multiple visits between May and July 2014.

| Site | Total infected | Total sampled | Prevalence | pH |
| --- | --- | --- | --- | --- |
| DP | 0 | 68 | 0.000 | 5.11 |
| R10 | 20 | 359 | 0.056 | 4.69 |
| RNew | 118 | 278 | 0.424 | 5.38 |
| R4 | 296 | 476 | 0.622 | 6.26 |

Palmate newt skin communities also exhibited clear variation in overall composition (beta diversity) by site and infection class within site (Fig. 4). Despite infected and uninfected newts in site Rnew possessing similar mean bacterial richness (Fig. 3C), they differed markedly in community composition at the Phylum level (Fig. 4). Uninfected newts from DP and Rnew appeared broadly similar in composition, containing lower proportions of Proteobacteria and higher proportions of Actinobacteriota compared to infected newts in Rnew and both infection classes in R4. R10 newts possessed a relatively unique microbiome signature comprising roughly 10% Verrucomicrobiota (Fig. 4).

**Figure 4.**
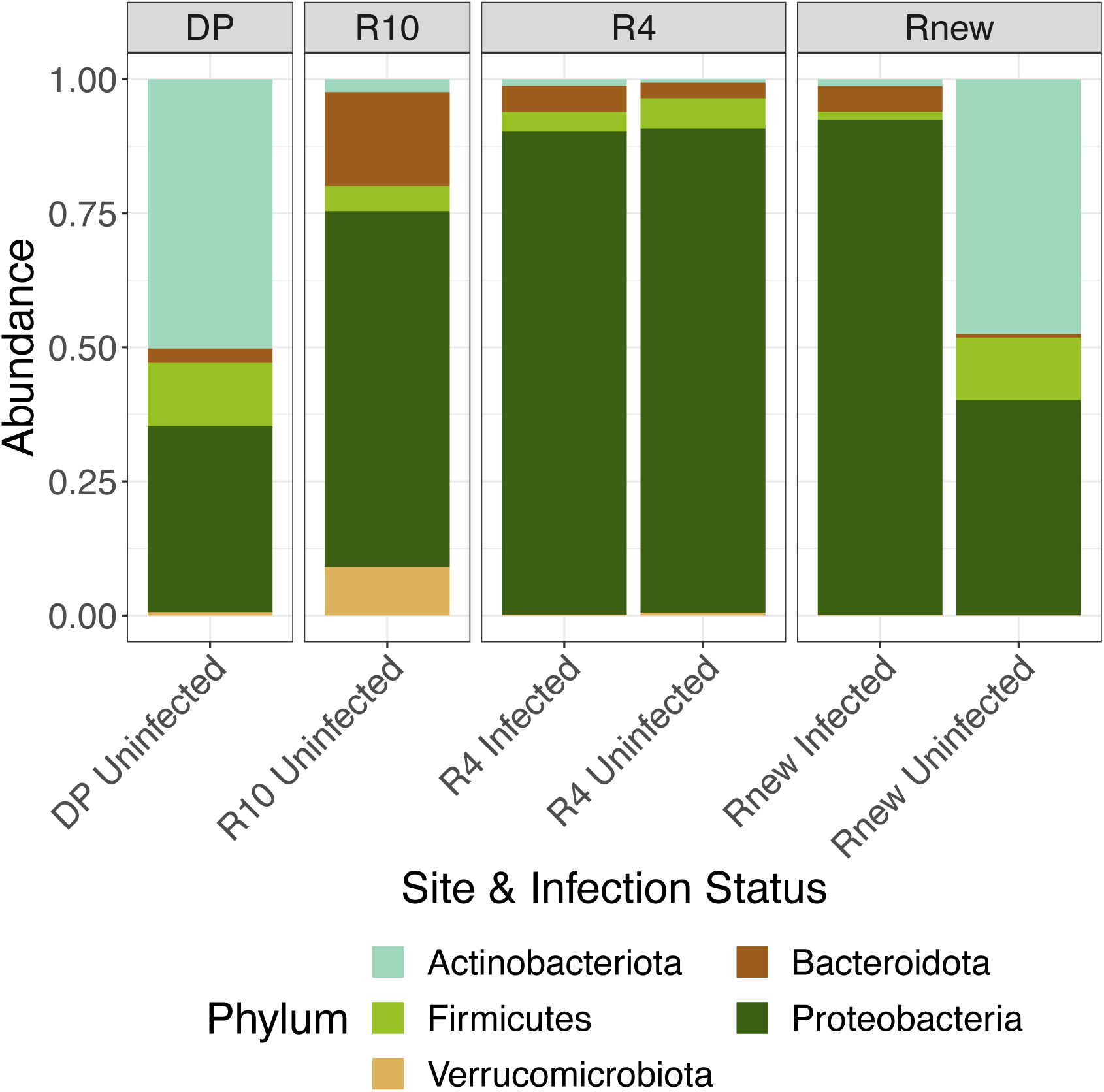
Variation in skin bacterial microbiota of palmate newts by site and infection class at the Phylum level.

Ordination of microbial community structures by site and infection class supported these patterns (Fig. 5A). PERMANOVA analysis indicated that both site (r2 = 26.6%) and infection class (r2 = 2.8%) were significant drivers of differences in bacterial composition (Table S5). Constrained Correspondence Analysis (CCA) similarly found both pH and infection class (p<0.001; Table S6) but not SVL (p=0.88) to influence skin microbiota structure (Fig. 5B). For both PCA and CCA ordinations, uninfected newts from Rnew appeared more similar to DP individuals, whilst infected individuals had microbial signatures that converged upon those of R4 newts (Fig 5A,B), consistent with the Phylum level patterns (Fig. 4).

**Figure 5.**
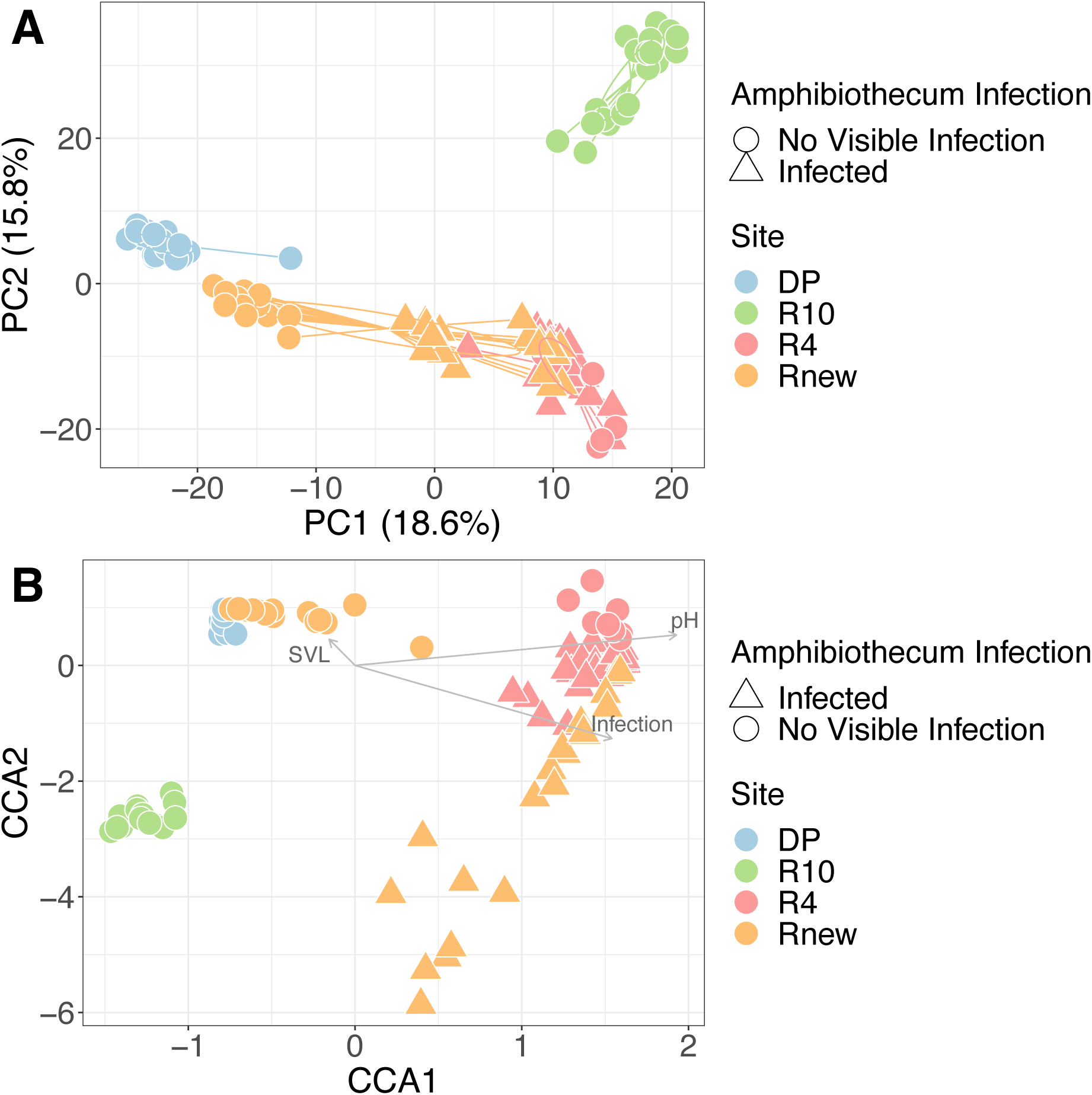
Community composition of Palmate newts sampled across 4 sites (point colours) and infection class (point shapes). **(A)** PCA ordination of Euclidean distances among CLR-transformed unrarefied community abundances **(B)** CCA ordination with biplot arrows that show loadings of model variables onto the 2 primary constrained axes – Infection and pH were both significant drivers of community composition, whereas SVL was not significant.

Differential abundance modelling using general linear latent variable models identified a significant site:infection interaction. Several bacterial genera differed between infected and uninfected newts, but these effects were site-specific within R4 and Rnew (Fig. 6A). Infected newts at R4 possessed significantly higher relative abundances of genera such as Pseudomonas and Parabacteroides, but this pattern was not mirrored in site Rnew. Instead, Rnew possessed a unique fingerprint of bacterial general associated with infection, including significant depletion of genera with potential protective effects such as *Paenibacillus*. Several genera with known anti-fungal properties showed a significant site:sex interaction, including *Chryseobacterium* and *Janthinobacterium* (Fig. 6B). Other genera like *Pseudomonas* showed site specific patterns, being at higher mean relative abundance in infected newts in site R4, but significantly lower in infected newts in Rnew. We also identified several blocks of bacterial genera that seemed to cooccur in newts after accounting for site and infection (Fig. S3). These included opportunistic pathogens such as *Pseudomonas*, *Stenotrophomonas* and *Klebsiella*, commonly found in freshwater environments.

**Figure 6.**
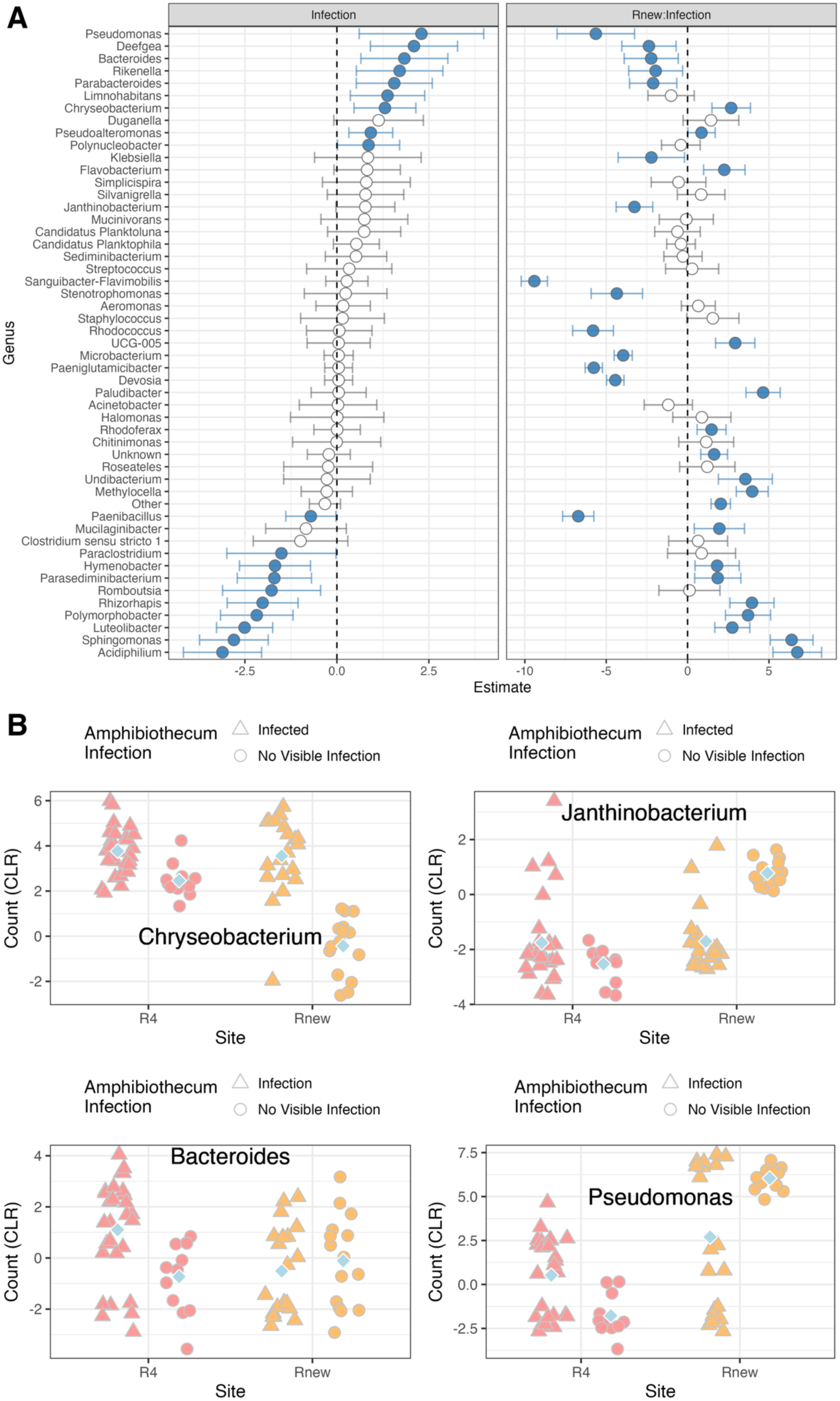
(A) Differential abundance of bacterial genera between infected and uninfected newts within sites for R4 and Rnew. Significant differences are denoted in blue for genera where 95% confidence estimates did not include zero **(B)** CLR-transformed abundances of key genera identified in the General Linear Latent Variable modelling in (A). Each point is the relative abundance of a genus in that sample grouped by site and infection status. Group means denoted by the blue diamond.

### Site-Specific Core Microbiota and Predicted Function

Sites varied markedly in their core microbiota, both in frequency of ASVs and composition (Fig. S3A). DP and R10, both more acidic ponds with low *Amphibiothecum* prevalence sites, had the most core ASVs (Fig. S4A) and the most unique core ASVs (Fig. S4B). The DP core microbiota contained several low-pH and halotolerant genera such as *Microbacterium* and *Devosia* (Table S8). Predicted functional analysis revealed significant variation in predicted metagenome by site (Fig. 7A) and infection class (Fig. 7B). We identified several KO terms that were enriched in uninfected newts (Fig. 7C), including those involved in pathways for Type IV secretion systems, the Ethylmalonyl pathway (essential to carbon fixation in Alphaproteobacteria), the methionine salvage pathway, and assimilatory nitrate reduction. A cluster image map revealed systematic separation between infected and uninfected newts in predicted metagenome profiles (Fig. 7D). The clustering analysis also identified a block of KO terms seemingly absent in >50% of infected newts, and present in higher abundance of most uninfected newts, which was linked to pathways involved in menaquinone synthesis and the two-component regulatory system (antimicrobial resistance and amino acid metabolism).

**Figure 7.**
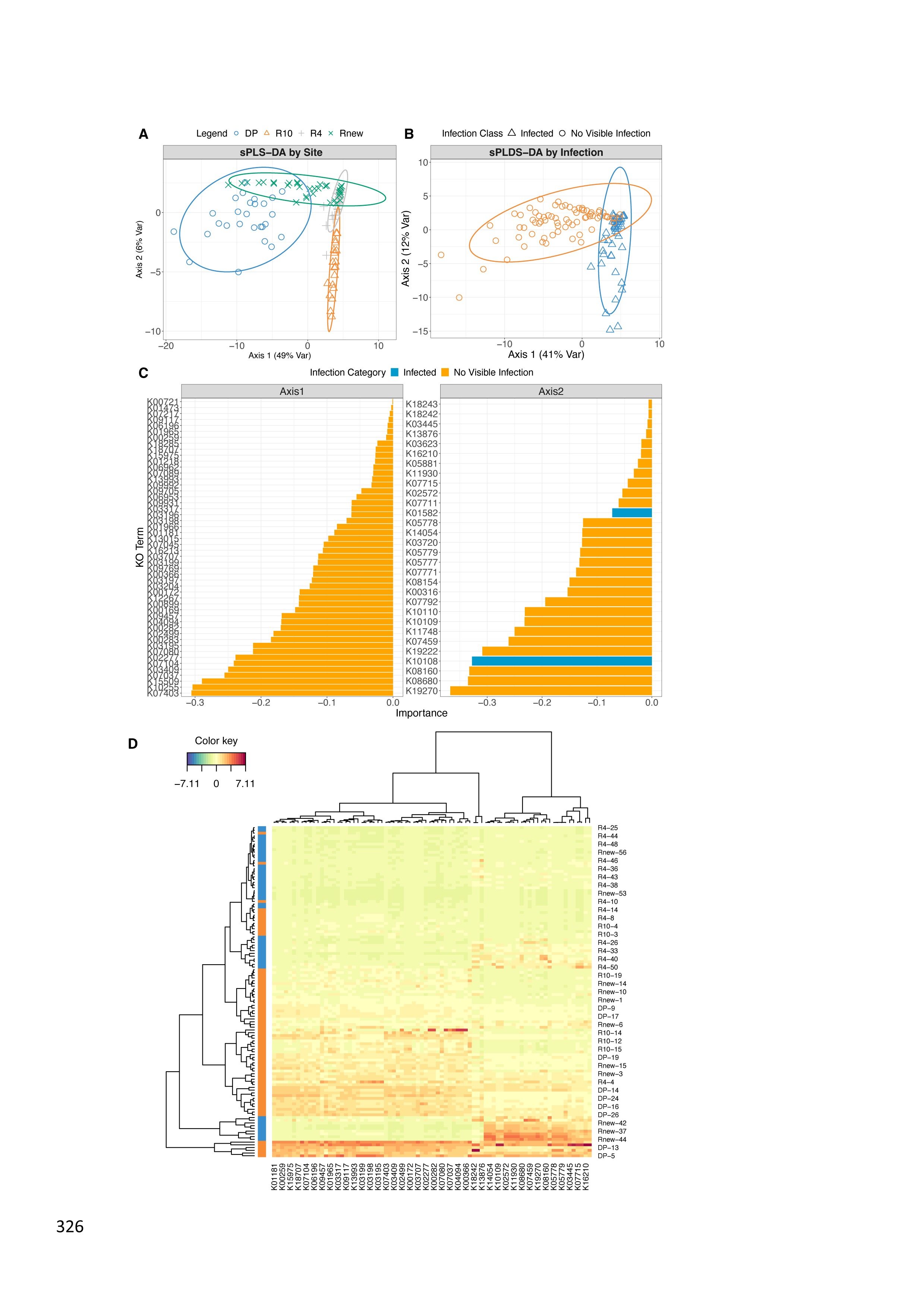
Predicted functional profiles of the newt skin bacterial microbiota. sPLS-DA analysis revealed significant variation in the functional metagenome by site **(A)** and infection class **(B)**, and identified a raft of KO terms that were differentially enriched in Uninfected newts **(C).** Cluster analysis tended to place uninfected newts together **(D),** with the exception of 3 infected newts from Rnew with low infection burdens, and identified large blocks of KO terms that appeared absent in most infected newts.

### Newt Skin Bacterial Communities Shift Over Time

We resampled 3 of the 2014 sites roughly 1 month after the initial surveys and detected significant shifts in skin bacterial community composition between visits (Fig. 8). PERMANOVA analysis identified a significant site:visit interaction explaining 7.7% of variation in bacterial community composition, in addition to a cumulative 28% of variation explained by the main effects of site and visit (Table S9).

**Figure 8.**
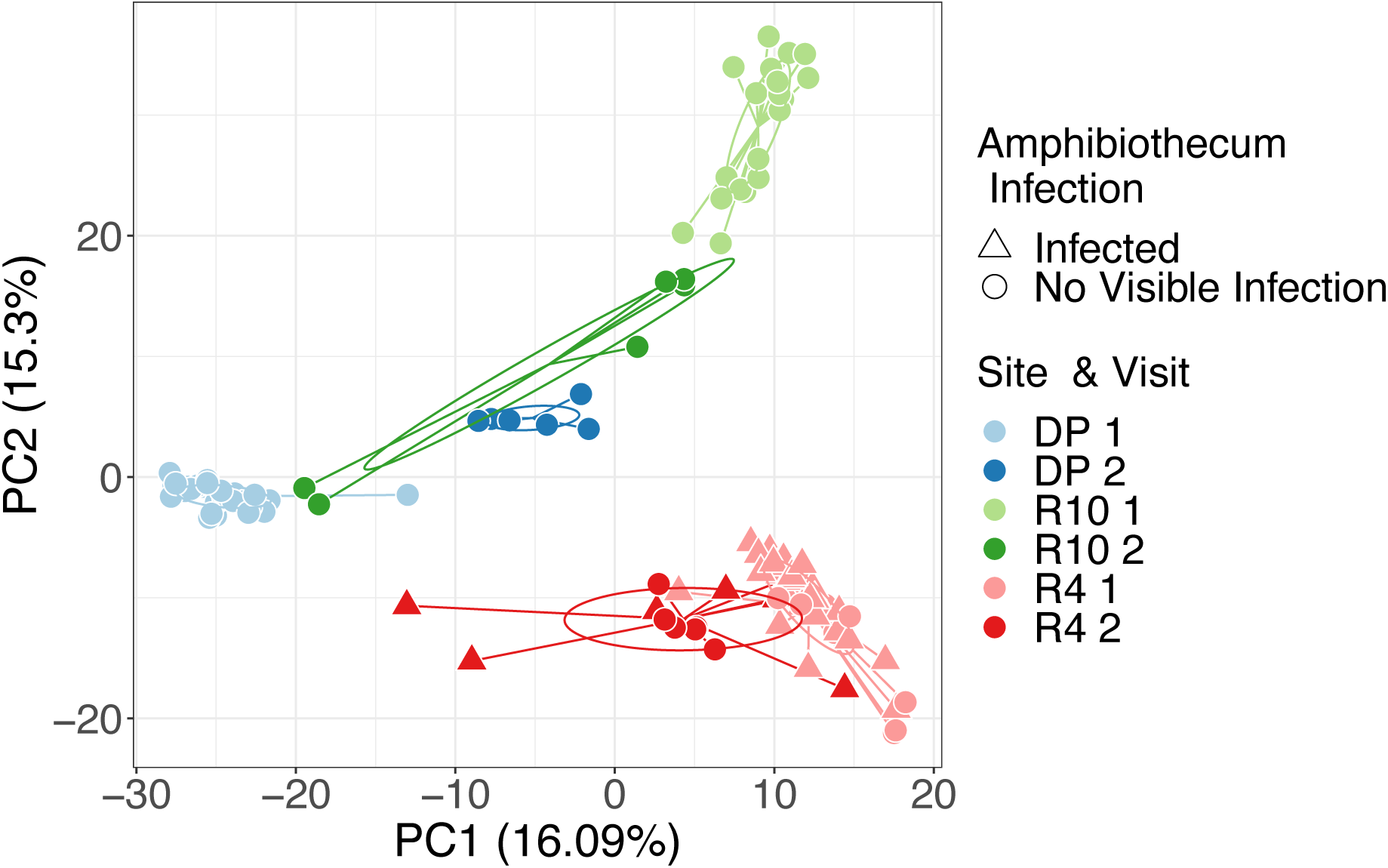
Temporal variation in the Palmate newt skin microbiome over time. Sites are shaded in paired colour palettes where darker shades represent the 2^nd^ visit (∼1 month later). Shape of points represents infection class (relevant for site R4).

## DISCUSSION

In this study we have identified an association between environmental pH, the dynamics of the Palmate newt skin microbiome, and prevalence of infection with the emerging pathogen *Amphibiothecum meredithae.* The distribution of disease was largely explained by environmental variation; sites where the pond pH was lowest and temperatures colder had consistently lower levels of infection with *A. meredithae*. These data are consistent with previous work on multiple host-pathogen systems that have identified a link between pH, temperature and infection prevalence e.g. protozoans (*Ichthyophthirius multifiliis*) infecting fish (Garcia et al., 2011), fish-infecting dermocystid parasites (reviewed in Gozlan et al., 2014) and the fungal pathogen *Bd* infecting amphibians (Turner et al., 2021). Studies of *Bd-*infected salamanders (Basanta et al., 2023), anurans (Turner et al., 2021) and the environmental density of *Bd* zoospores (Blooi et al., 2017) have invoked acidity as an environmental factor that reduces zoospore density and prevalence. In our system, we hypothesize that low pH reduces endospore survival in the environment which in turn reduces the risks of transmission. Furthermore, although the propagation of *A. meredithae* among host tissues has been described, increased encysting in the skin is more likely driven by reinfections (Fiegna et al., 2017). If this is the case, reduced pH may also serve to limit the escalation of the severity of infections, which in turn should decrease mortality.

*Infection and pH Shape Palmate Newt Skin Microbiome Composition and Predicted Function* We found a strong association between environmental pH and the bacterial richness of the Palmate newt skin microbiome. Bacterial alpha diversity increased across the four ponds in a non-linear fashion as pH decreased; previous work on tropical dendrobatid frog species has also reported a similar relationship where skin microbiotas are more diverse in more acidic soil (Varela et al., 2018). Associations between pH and microbiome also correlated with infection states, where more acidic ponds (e.g. R10) exhibited no evidence of visible infection in our sample. While site R4 exhibited microbiota diversity comparable to site DP, this was largely driven by the effect of uninfected newts in R4, which carried significantly greater diversity than the infected animals at the same location (Fig 3B). The effect of site and infection status on bacterial community composition was further supported through ordination (Fig. 5), suggesting a combined relationship between low pH, increased microbial diversity and the presence and prevalence of *A. meredithae.* Longitudinal sampling of bacterial microbiotas revealed that distinct signatures of populations and infection status were maintained over time (Fig. 8), indicating the ‘site-level’ signature is relatively stable and adding further support for the relationship between location, microbiome community structure and disease. Previous work on wild *Craugastor ranoides* in Costa Rica revealed distinctive signals of ‘individuality’ in the skin microbiome even after translocation (Harrison et al., 2025), suggesting that stability of the microbiome at the individual level may constrain the extent of population-level shifts over time.

Increased microbial diversity is hypothesized to be a response to or a bet hedge against amphibian infectious diseases, in that greater diversity is more likely to contain constituents that are actively antiparasitic (Bates et al., 2018; Federici et al., 2015; Harrison et al., 2019; Muletz Wolz et al., 2018; Muletz-Wolz et al., 2017). In support of this, we found evidence of differences in functional profiles of the Rum Palmate newt microbiomes that were largely distinct based on infection status (Fig. 7), even though both overall taxonomic and core microbiome profiles were inconsistent across sites (Fig. 4, Fig. 6). These functional differences were the result of enriched pathways in uninfected animals, including mechanisms underpinning antimicrobial resistance such as the bacterial Two-Component System (TCS) (Bates et al., 2022). Antimicrobial resistance afforded by TCS genes likely permit beneficial skin-associated microbes to subvert these defences to maintain mutualistic interactions with the host (Bates et al., 2022). It is worth noting that enrichment of antimicrobial mechanisms also correlated with increased bacterial diversity, indicating that antimicrobial activity is not associated with deleterious disruption of the microbiome (Jani et al., 2021). Based on our correlations, they suggest a functional response to the risk of infection (Bates et al., 2022; Rebollar et al., 2018).

We also found other pathways linked to the TCS were enriched in uninfected newts. The TCS plays an important role in responding to environmental stimuli, including pH (reviewed in Xie et al., 2022) and may play a role regulating bacterial persistence on amphibian skin exposed to low pH pond water (Tiwari et al., 2017). Furthermore, we found predicted gene pathways linked to the Type IV secretion system (T4SS). Type IV secretion systems perform a variety of functions, including biofilm formation (T. R. Costa et al., 2024) host colonisation, and bacterial competition (Souza et al., 2015). Variation in the patterns of T4SS pathways may reflect heightened competition for ecological niches on healthy newt skin, niches that are compromised by the damage done to skin by infection with *A. meredithae*.

Collectively our data suggest that lower pH is hostile for *A. meredithae* endospore survival, impairing transmission and reinfection rates. At the same time, lower pH allows newts to support a more diverse host skin microbiome that is functionally reinforced by that community and possibly inhibitory to *A. meredithae infections*. Previous work has shown increases in bacterial richness of the skin microbiota with more acidic pH in *Bombina variegata* (Lucia et al., 2025), and covariation of skin microbiome with environmental pH in *Pelophylax perezi* (S. Costa et al., 2016). Similarly, drought-mediated shifts in the microbiome have been shown to increase susceptibility to the fungal pathogen *Bd* (Buttimer et al., 2024). Taken in tandem with our data, these studies suggest that shifts in the microbiota composition driven by the abiotic environment may be a general mechanism explaining variation in patterns of infection in wild amphibians. At the landscape scale, heterogeneity in pH among ponds mean that palmate newts on Rum can occupy and breed in disease-free ponds despite proximity to, and likely immigration from, ponds where newts are dying from disease caused by *A. meredithae*. Why, then, don’t Palmate newts avoid less acidic and neutral pH sites where the risk of infection is markedly higher? We propose two mechanisms. First, infection and disease are not absolute at high-risk locations so breeding by healthy newts can still be supported at locations where disease occurs. Second, pH values where disease does abundantly occur have been previously shown to impair larval growth and development (Griffiths et al., 1993). Amphibian reproductive strategies are commonly structured by trade- offs in oviposition site choice and subsequent costs for larval development (Goldberg et al., 2006; Gould et al., 2021; Van Buskirk, 2002). Ultimately, trade-offs are driven by optimizing recruitment (Orizaola et al., 2013), not adult survival, so a Palmate newt reproductive strategy that involves increased risk of adult mortality to the benefit of larval survival through to metamorphosis would be adaptive. Whatever the mechanisms are that supports widespread newt exploitation of ponds on Rum, though, our study shows that potentially lethal, single host species parasites and their host can coexist within highly constrained geographical ranges through processes largely governed by environmental heterogeneity. Furthermore, small-scale environmental heterogeneity can support hosts through the maintenance of beneficial microbial communities that generate beneficial host microbiomes. Amphibians are beset by declines driven by infectious disease and many of those worst affected are small range size species (Luedtke et al., 2023; Scheele, Pasmans, et al., 2019). The scale of the problem precludes eradication as a conservation strategy, while managing landscapes for host and parasite coexistence has been proposed as a more effective strategy (Scheele, Foster, et al., 2019). Our data support this proposition.

## Supporting information

Supplemental Figutres

## ACKNOWLEDGEMENTS

XAH acknowledges financial support from the Leverhulme Trust (RPG-2020-320).

## DATA AVAILABILITY

Raw sequence data have been deposited at the Sequence Read Archive (SRA) under project ID PRJNA1294852. All code and datasets to reproduce the analyses in this work are lodged in a GitHub repository at https://github.com/xavharrison/NewtsOnRum2024

