## Supplemental Figutres for "Environment shapes the association between the skin microbiome and infection dynamics of the Dermocystid pathogen *Amphibiothecum* in Palmate newts (*Lissotriton helveticus*)"

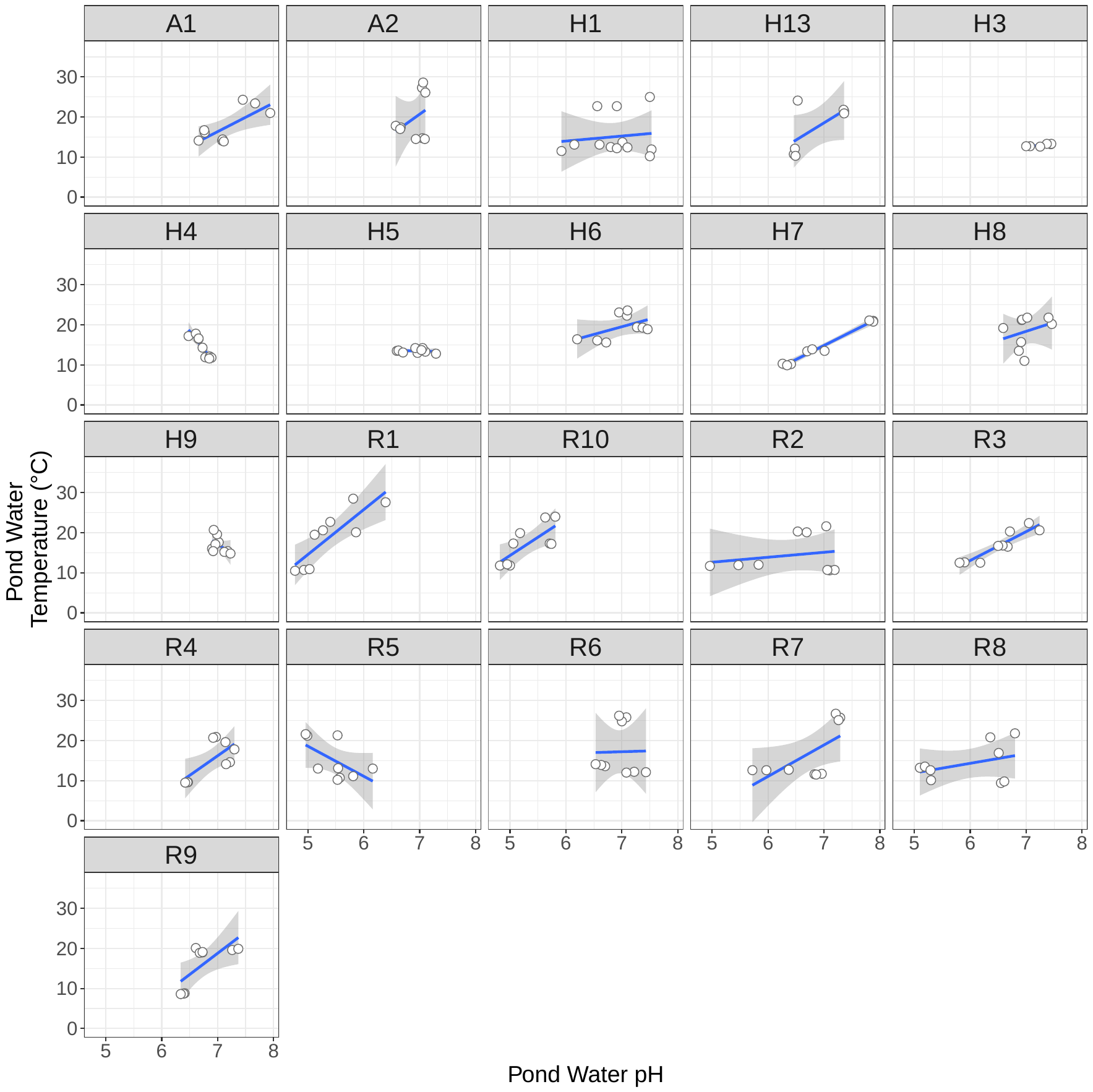

**Figure S1. Relationship between pond temperature and pH for 21 sites (n=187 measurements).** A Bayesian mixed model controlling for site (random intercept) and day of measurement (day of year) recovered a positive association between temperature and pH (Table S1A). A random slope equivalent allowing slope to vary by site did not converge.

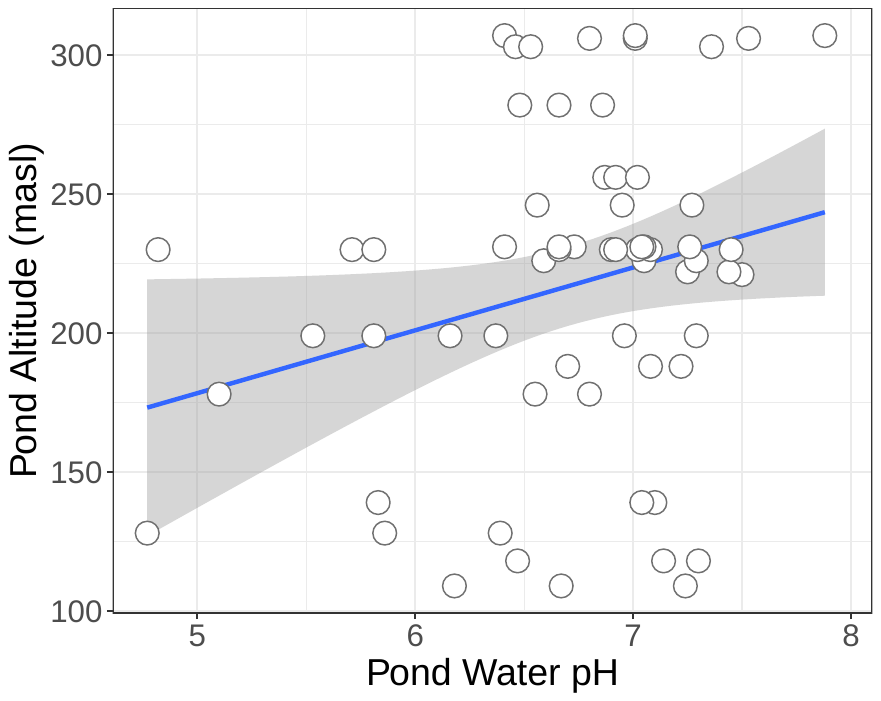

**Figure S2. Relationship between pond altitude and pH for 21 sites.** A Bayesian mixed model controlling for site (random intercept) found no association between altitude and pH (Table S1B)

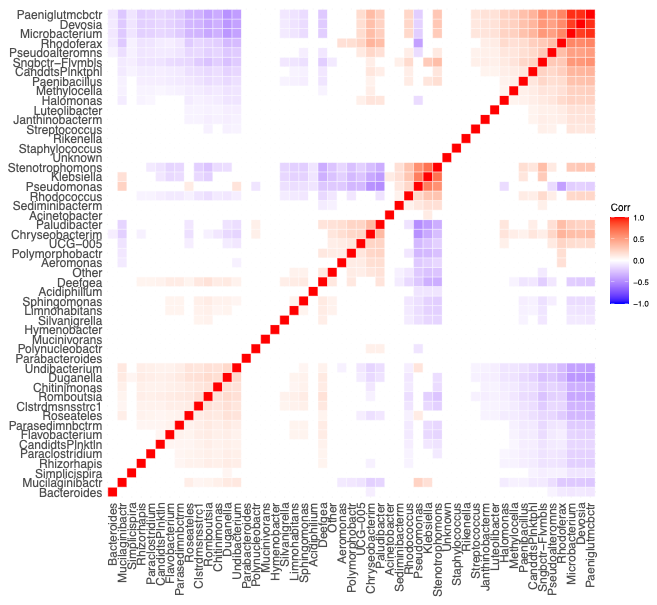

**Figure S3.** Residual correlation plot of a general linear latent variable model after accounting for site and infection effects, highlight positively (red) and negatively (blue) associated sets of bacterial genera.

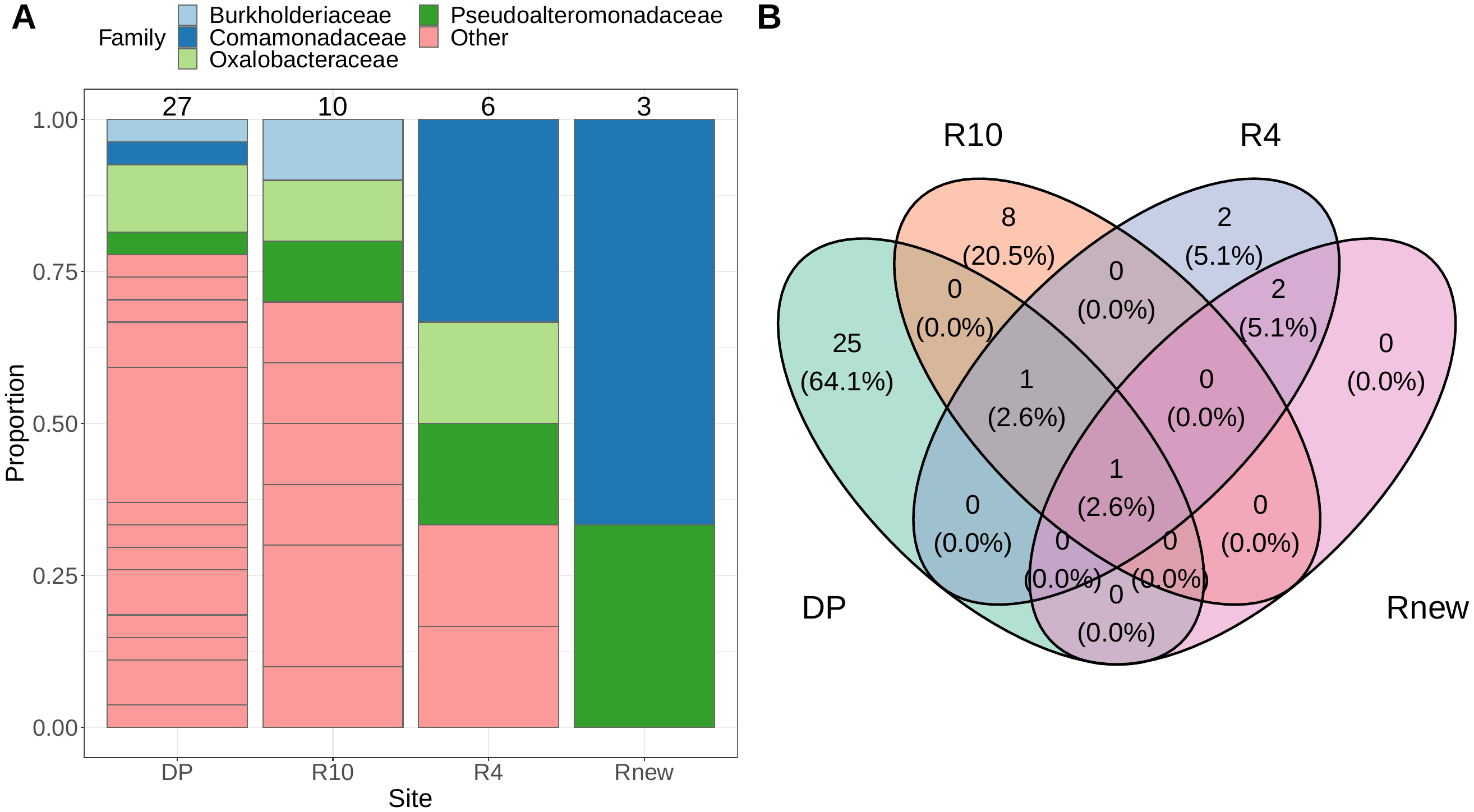

**Figure S4.** Core microbiota variation by site, where core ASVs are identified as occurring in at least 80% of samples from a site **(A)** Family-level taxonomy of ASVs occurring in at least 80% of samples within a site, with total number of core ASVs shown above bars. Taxonomy is grouped at the Family level. Families with only a single ASV have been collapsed into the ‘Other’ category for clarity of plotting **(B)** Venn diagram showing number of unique and shared core microbiota ASVs by site.

| **A) pH and Pond Temperature** | | | | | | | |
| --- | --- | --- | --- | --- | --- | --- | --- |
| ~site (Number of levels: 21) |  |  |  |  |  |  |  |
|  | **Estimate** | **Error** | **Lower 95%** | **Upper 95%** | **Rhat** | **Bulk_ESS** | **Tail_ESS** |
| sd(Intercept) | 0.61 | 0.11 | 0.43 | 0.87 | 1 | 624 | 1093 |
| Population-Level Effects: |  |  |  |  |  |  |  |
|  | **Estimate** | **Error** | **Lower 95%** | **Upper 95%** | **Rhat** | **Bulk_ESS** | **Tail_ESS** |
| Intercept | 5.56 | 0.22 | 5.11 | 5.99 | 1 | 791 | 1327 |
| Pond Temperature (C) | 0.07 | 0.01 | 0.05 | 0.09 | 1 | 1757 | 2448 |
| Date (DOY) | -0.15 | 0.05 | -0.25 | -0.05 | 1 | 1631 | 2182 |
| Family Specific Parameters: |  |  |  |  |  |  |  |
|  | **Estimate** | **Error** | **Lower 95%** | **Upper 95%** | **Rhat** | **Bulk_ESS** | **Tail_ESS** |
| sigma | 0.4 | 0.02 | 0.36 | 0.45 | 1 | 2552 | 2744 |
| **B) pH and Altitude** | | | | | | | |
| ~site (Number of levels: 21) |  |  |  |  |  |  |  |
|  | **Estimate** | **Error** | **Lower 95%** | **Upper 95%** | **Rhat** | **Bulk_ESS** | **Tail_ESS** |
| sd(Intercept) | 0.44 | 0.12 | 0.21 | 0.7 | 1 | 1004 | 1151 |
| Population-Level Effects: |  |  |  |  |  |  |  |
|  | **Estimate** | **Error** | **Lower 95%** | **Upper 95%** | **Rhat** | **Bulk_ESS** | **Tail_ESS** |
| Intercept | 6.19 | 0.45 | 5.3 | 7.11 | 1 | 2188 | 2844 |
| Alt | 0 | 0 | -0.01 | 0.01 | 1 | 2290 | 2831 |
| date_doy_z | 0.1 | 0.07 | -0.03 | 0.23 | 1 | 5459 | 2717 |
| Family Specific Parameters: |  |  |  |  |  |  |  |
|  | **Estimate** | **Error** | **Lower 95%** | **Upper 95%** | **Rhat** | **Bulk_ESS** | **Tail_ESS** |
| sigma | 0.49 | 0.06 | 0.4 | 0.62 | 1 | 1983 | 2029 |

**Table S1. Model output for relationship between pH and A) temperature and B) altitude.** Model is a Bayesian GLMM with random site intercepts. A random slope model with varying slopes for temperature/altitude given site would not converge.
